# A Data-Driven Model to Identify Fatigue Level Based on the Motion Data from a Smartphone

**DOI:** 10.1101/796854

**Authors:** Swapnali Karvekar, Masoud Abdollahi, Ehsan Rashedi

**Affiliations:** Biomechanics and Ergonomics Lab, Industrial and Systems Engineering Department Rochester Institute of Technology, Rochester, NY, US

**Keywords:** Fatigue detection, Gait analysis, Machine learning, Smartphone, Wearable technology

## Abstract

The fatigue due to repetitive and physically challenging jobs may result in workers’ poor performance and Work-related Musculoskeletal Disorder (WMSD). Thus, it is imperative to frequently monitor fatigue and take necessary recovery actions. Our purpose was to develop a methodology to objectively classify subjects’ fatigue level in the workplace utilizing the motion sensors embedded in the smartphones. An experiment consisting of twenty-four participants (12 M, 12 F) with a smartphone attached to their right shank was conducted using a fatiguing exercise (squatting), targeted mainly the lower extremity musculature. After each set of an exercise (2-min squatting), participants were asked about their ratings of perceived exertion (RPE), then a reference gait data were collected during a straight walk of 20-32 steps. This process was continued until they reported strong fatigue (≥17). Using the RPE to label the gait data, we have developed machine learning algorithms (i.e., binary and multi-class SVM models) to classify the individuals’ gait into two (no-vs. strong-fatigue) and four levels (no-, low-, medium-, and strong-fatigue). The models reached the accuracies of 91% and 61% for two and four-level classification, respectively. The outcomes of this study may facilitate the implementation of a proactive approach in continuous monitoring of operators’ fatigue level, which may subsequently increase the workers’ performance and reduce the risk of WMSDs.

## 1. Introduction

Human muscle fatigue has been known to negatively impact the performance and linked to the risk of WMSDs [1]. One out of every three dollars spent on workers compensation is related to a WMSD claim, according to the Occupational Safety and Health Administration [2]. In addition, employers spend over $20 billion a year on direct WMSD-related compensation, demonstrating the importance of reducing WMSD incidents [2].

A number of studies have been conducted to detect fatigue with the goal of reducing the risk of developing WMSDs. Among these, some have used wearable devices for measuring kinematic and kinetic parameters of human gait. For example, the inertial measurement unit (IMU) systems have gained attention for gait analysis, since the sensors are small, light and portable, thus are minimally intrusive for a range of applications [3, 4]. Recently, few researchers have used similar technologies to quantify human muscle fatigue. In a study to estimate fatigue during running, Buckley et al. placed one IMU on each of the lumbar spine and both shanks [5]. They collected the IMU data before and after the fatiguing exercise and utilized the Borg’s RPE of ≥18 as the fatigued level, where they have achieved a maximum accuracy of 77% in identifying fatigue. In another recent study investigating fatigue-induced gait changes during an occupational task, Baghdadi et al. employed a single IMU strapped at the right ankle of participants [6]. Specifically, the fatiguing exercise consisted of three-hour manual material handling tasks, where participants transported weight containers. Authors reported 90% accuracy in differentiating between the high level of fatigue and no-fatigue conditions [6].

While achieving convincing results, the studies noted above were limited to the laboratory environment and their specific equipment. The next imperative step can be to investigate methods to improve the accessibility of fatigue assessment in the workplace. Interestingly, similar IMU technology is also embedded in today’s smartphones. With the widespread access to these devices, we may use them to identify different levels of fatigue in an industrial environment. Besides the availability, smartphones are ubiquitous, easy to carry, feature a range of applications, and can be connected to a number of devices through Wi-Fi and Bluetooth, which may substantially enhance the capability of fatigue assessment in the workplace. Hence, this study aimed to develop a model to identify the fatigue level based on the smartphone signal of walking, as walking is a prevalent activity in the workplace. A smartphone was attached to the shank of the participants, and as they walked, IMU data was collected by smartphone. This IMU data was labeled according to the participant’s rating of RPE, and then binary/multi-class SVM was used to classify the IMU data into different levels of fatigue.

## 2. Methodology

### 2.1 Participants

Twenty-four participants (12M, 12F) were recruited according to the following acceptance criteria: (a) must show no current or recent history of musculoskeletal disorders or lower-body injury, (b) must be engaged in exercise 2-3 days per week, and (c) must be between 18 and 35 years of age. The participants were recruited from the student population at the Rochester Institute of Technology (RIT). Prior to data collection, informed consent (as mandated by RIT’s IRB) was collected from all participants.

### 2.2 Experimental Procedure

Borg’s ratings of perceived exertion is one of the frequently used subjective measures to assess the exertion level [5, 6], which combines the feelings of physical stress, and fatigue [7]. To reduce the inherent errors in this subjective measure, participants’ perception of exertion was first calibrated by conducting the knee test similar to our earlier studies, where participants were asked to lean against the wall with their knees bent at 90° until their RPE reached a value of ≥18 [8, 9]. Prior to the exercise phase, a smartphone was attached to participants’ shank using Velcro straps as shown in figure 1. The position of smartphone at shank was chosen after performing a pilot study with accessing smartphone’s positions at different locations such as shank, thigh and considering the accessibility of smartphone to workers. After sensor placement, data was collected during a walking trial comprising of 20-32 steps, followed by sets of 16 squats (8 squats/min). Upon completion of each squat set, participants were asked to state their Borg’s RPE rating and then walked again for 20-32 steps. The IMU data was labeled according to the participant’s Borg’s rating. These cycles of walking/squatting continued till the Borg’s rating was ≥ 17.

**Figure 1:**
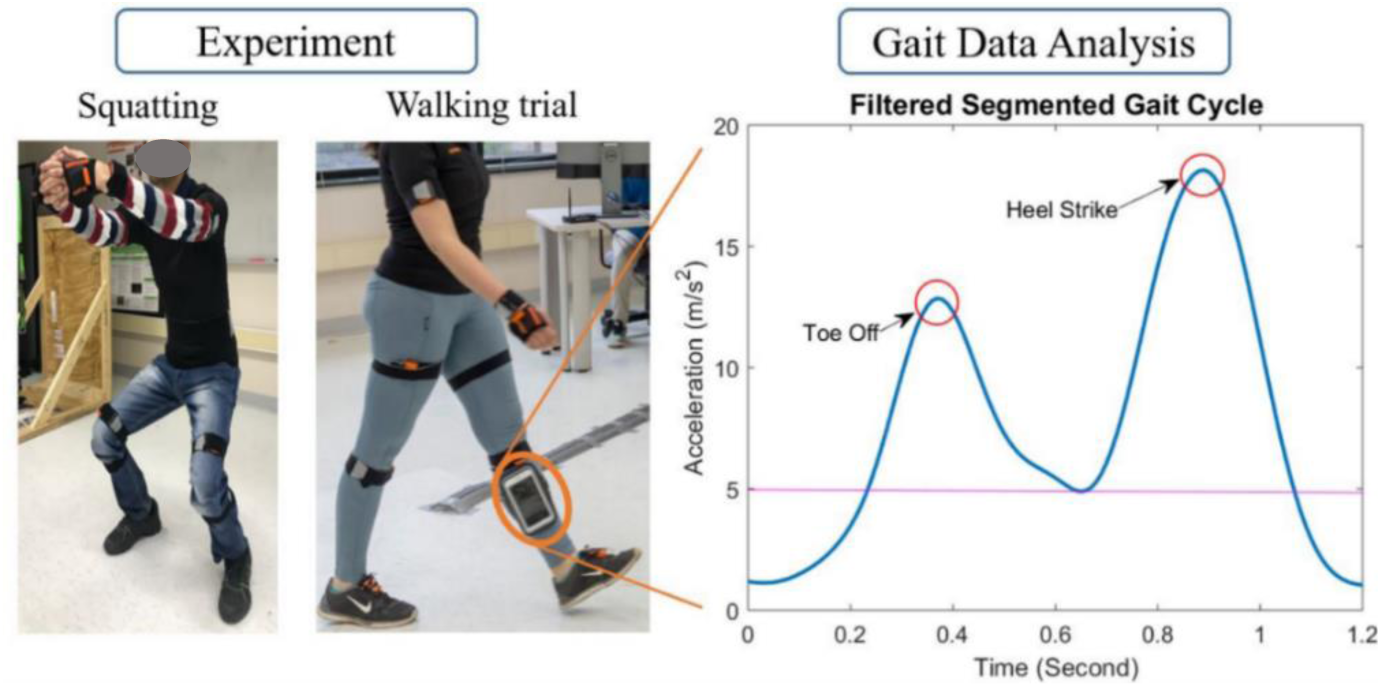
Schematic system diagram of the platform for experimental procedure and data analysis

### 2.3 Instrumentation

IMU data was collected from the smartphone at 100Hz. We conducted one pilot study to study the performance of IMU embedded in the smartphone and the Xsens system. Specifically, the smartphone was attached to the participants’ upper right leg along with the Xsens system [10]. The captured data from both equipments were synchronized by asking the participants to jump at the start of each walking trial. Figure 2 shows the overlapping signals from the Xsens sensor (in blue) and the Smartphone (in orange). It can be seen that both signals show good conformity as the correlation coefficient is more than 0.8.

**Figure 2.**
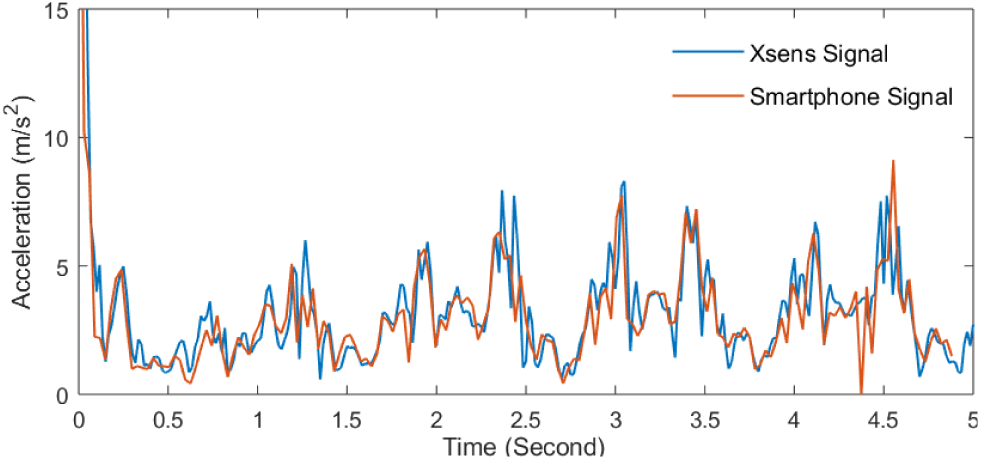
IMU Data from Xsens sensor and Smartphone

### 2.4 Data analysis

The IMU data collected from the smartphone was processed to get segmented gait cycle after filtering it by Butterworth filter (order 4 and a cutoff frequency of 3 Hz) as generally, the walking frequency of a human being is 0.6 Hz to 2 Hz [11]. From this segmented gait cycle, some features were extracted, which were then used to classify gait cycles into different fatigue levels.

#### 2.4.1 Gait segmentation

In a sliding window algorithm, the length of the window is fixed, and window slides along continuous IMU data points to identify small segments of gait cycles. In this study, the sliding window of 70-80 acceleration data points was utilized. As shown in Figure 1, the gait cycle for one leg consists of two peaks, which is higher for heel-strike and smaller for toe-off [12]. As a normal person requires 1-1.2 seconds to complete one gait cycle and the larger peak and start of cycle constitute approximately 75% of the cycle, the sliding window of 70-80 points was used. The data analysis was performed in MATLAB R2018a (MathWorks, Inc., Natick, MA, USA).

If for any window the ratio of larger peak to smaller peak is greater than 1.1, with global maximum higher than 5 m/s^2^, then that window was considered to correspond to a valid gait cycle [6], as shown in equation (1).

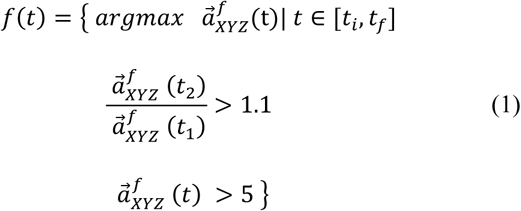

Where *t*_*i*_ and *t*_*f*_ are the times at which gait cycle starts and ends. *t*_1_and *t*_2_ are the times at which a smaller and larger peak occurs, and 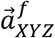 is the filtered resultant acceleration. The end of the gait cycle was considered as the minimum value of acceleration from the start of the larger peak and 50 points after that. The data points from the start point of the sliding window to the minimum value of acceleration were considered as a one gait cycle. The next sliding window starts from end of the previous cycle.

After gait segmentation, various features including mean, coefficient of variation, maximum acceleration, acceleration range, second acceleration peak, skewness, kurtosis, minimum and maximum vertical acceleration, minimum and maximum frontal acceleration, area under curve etc. [13–15] were extracted from each gait cycle, which were selected from the literature.

#### 2.4.2 Fatigue Classification

The mentioned features were used as input for the one-versus-one multi-class SVM classifier with Radial Basis Function kernel. The purpose of this study was not to predict the exact values of fatigue as a continuous variable, rather some general notion of fatigue was satisfactory, i.e., no and strong fatigue (2 classes), or no, low, medium, and high fatigue levels (4 classes). While the 2-class SVM can be helpful to identify the high levels of fatigue, in real life, we do not want the workers to experience very large fatigue levels (e.g., RPE ≥17). It would be helpful to identify for example medium (RPE of 11-15) to high-levels of fatigue (RPE ≥15). Therefore, we also incorporated the 4-class SVM besides the 2-class analysis (Table 1).

**Table 1:** Participant’s fatigue classification based on the measure of ratings of perceived exertion (RPE)

|  | Normal Walking | 7-10 | 11-15 | 15-20 |
| --- | --- | --- | --- | --- |
| **2-class SVM** | 0 | NA | NA | 1 |
| **4-class SVM** | 0 | 1 | 2 | 3 |

## 3. Results and Discussion

To enhance the accessibility of fatigue detection, this study was aimed to explore the possibility of identifying different human muscle fatigue levels using smartphones. An experiment was conducted to collect the motion data from a smartphone during the participants’ walk. Classification accuracy of 91% and 61% were achieved for 2-class and 4-class SVM, respectively.

Use of gait analysis for identifying fatigue level has been studied in prior studies [5, 6, 15]. Buckley et al. were able to classify high-level fatigue using IMU attached to each of the lumbar spine and both shanks with a maximum of 77% accuracy [5]. However, they utilized running tasks for fatiguing the athletes, which had a fast pace fatiguing protocol with limited applicability in an industrial environment. In another study, Baghdadi et al. have investigated the capability of using a single IMU attached to the ankle to classify fatigue state [6] and reported the accuracy of 90% using IMU data from 20 participants. The fatiguing exercise consisted of three-hour manual material handling tasks, where participants transported weight containers. In comparison to our study, the main differences are the position of IMU sensor, the fatiguing protocol, and the features used. Similarly, in our study, we were able to identify the fatigued levels using smartphone with almost the same accuracy (i.e., 91% for 2-class).

As workers continue to perform in the high-level fatigue stage (RPE≥15), it not only increases risk of getting injured and developing WMSDs but also affects their performance and concentration resulting into poor quality of work. To address this concern, we tried to identify low-, medium-, high-level of fatigue (61% accuracy), which may help in monitoring fatigue levels at earlier stages to potentially implement a range of interventions including administrative controls (e.g., job rotation or changing the task parameters [9]) or engineering controls (e.g. task design [16], or using the assistive devices [8]).

With the help of features such as coefficient of variation and acceleration range, the accuracy of 91% was achieved for 2-class SVM. Figure 3 shows the confusion matrix for this model. The algorithm has 92% sensitivity and 90% specificity. In fact, identifying a no-fatigued person as a fatigued one, i.e., 63 cases in confusion matrix would not lead to a high-risk situation. However, if a fatigued person is identified as a no-fatigued one, i.e., 47 cases in confusion matrix it would increase the risk of injury. Subsequently, the model is prone to classify the workers more inside the safe margins (63 vs. 47) even in cases that there is confusion between the two classes.

**Figure 3:**
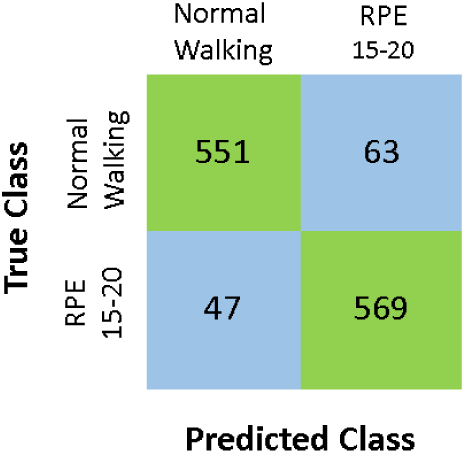
Confusion matrix for 2-class SVM

Accuracy decreased from 2-class SVM to 4-class SVM, since for the 2-class SVM the two extreme fatigue levels were classified, i.e., no vs. strong fatigue with a large gap in between; whereas for the 4-class SVM, the entire range of the Borg’s scale was divided into 4 classes, i.e., there is no gap between the classes (Table 1). Although the accuracy decreased by increasing the number of classes, it is critical to notice and intervene at earlier stages of fatigue, which in turn might help in reducing injuries and WMSDs. Figure 4 shows the confusion matrix for 4 class SVM. The highest confusion was between 3 and 4 classes since the algorithm classified 124 class 3 data points as class 4 and 144 class 4 data points as class 3. Future data analysis may enhance the accuracy of our model by conducting a 3-class analysis, changing the boundaries for the current 4-classes, or further identifying new features.

**Figure 4:**
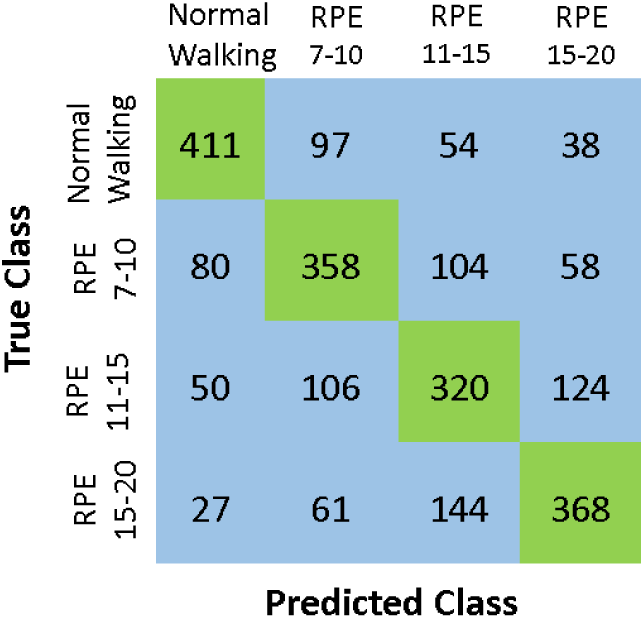
Confusion matrix for 4-class SVM

One limitation of this study was on the generalizability of the outcome. In fact, the participants were recruited from the university population, which may have different endurance limit and physical strength in comparison to workers. Besides, the fatiguing protocol was squatting exercise. Although this exercise involves many muscles same as that involved in the workplace activities, it is not an industrial task. Recruiting workers and industrial tasks to fatigue individuals can be considered in future study. Along with this, instead of subjective measure of fatigue, i.e. Borg’s rating of RPE objective physiological measures such as Maximum Voluntary Contraction (MVC) may be utilized to define various levels of fatigue. Despite these limitations, this study provides a unique approach to classify human muscle fatigue. The outcome of this study is enabling companies to take a proactive approach in continuous monitoring of operators’ fatigue level in their working environment without interfering with their daily activities, which in turn can reduce the risk of WMSDs and increase workers’ performance.

